# RetiGene, a comprehensive gene atlas for inherited retinal diseases (IRDs)

**DOI:** 10.1101/2025.06.08.653722

**Authors:** Mathieu Quinodoz, Elifnaz Celik, Dhryata Kamdar, Francesca Cancellieri, Karolina Kaminska, Mukhtar Ullah, Pilar Barberán-Martínez, Manon Bouckaert, Marta Cortón, Emma Delanote, Lidia Fernández-Caballero, Gema García García, Lara K Holtes, Marianthi Karali, Irma Lopez, Virginie G Peter, Nina Schneider, Lieselot Vincke, Carmen Ayuso, Sandro Banfi, Beatrice Bocquet, Frauke Coppieters, Frans P M Cremers, Chris F Inglehearn, Takeshi Iwata, Vasiliki Kalatzis, Robert K Koenekoop, José M Millán, Dror Sharon, Carmel Toomes, Carlo Rivolta

## Abstract

Inherited retinal diseases (IRDs) are rare disorders, typically presenting as Mendelian traits, that result in stationary or progressive visual impairment. They are characterized by extensive genetic heterogeneity, possibly the highest among all human genetic diseases, as well as diverse inheritance patterns. Despite advances in gene discovery, limited understanding of gene function and challenges in accurately interpreting variants continue to hinder both molecular diagnosis and genetic research in IRDs. One key problem is the absence of a comprehensive and widely accepted catalogue of disease genes, which would ensure consistent genetic testing and reliable molecular diagnoses. With the rapid pace of IRD gene discovery, gene catalogues require frequent validation and updates to remain clinically and scientifically useful. To address these gaps, we developed RetiGene, an expert-curated gene atlas that integrates variant data, bulk and single-cell RNA sequencing, and functional annotations. Through the integration of diverse data sources, RetiGene supports candidate gene prioritization, functional studies, and therapeutic development in IRDs.

## Introduction

The retina is a photosensitive tissue lining the posterior part of the eye. Its primary function is to convert light into electrical signals, which are then transmitted to the brain to form visual images. The retina contains two types of photoreceptors: rods and cones. Rods are responsible for vision in low light conditions, while cones provide sharp central vision, enable color perception, and mediate sight in bright light environments.^1^ The retinal pigment epithelium (RPE), a layer of pigmented cells located between the photoreceptors and the choroid, plays a crucial role in supporting vision. It absorbs excess light, forms part of the blood-retina barrier, transports nutrients and waste, regulates the visual cycle and removes photoreceptor debris, thereby ensuring their proper function.^2,3^ Other retinal cell types include retinal ganglion cells, horizontal cells, amacrine cells, Müller cells, among others, whose function is to encode visual signals detected by photoreceptors as well as to ensure the correct homeostasis of the retina by providing structural, metabolic, and immunological support.^4–7^

Inherited retinal diseases (IRDs) are a diverse group of monogenic conditions that typically lead to the progressive degeneration or dysfunction of photoreceptors, RPE cells, or other retinal neurons, culminating in vision loss and, in many cases, blindness. Clinically, IRDs are categorized based on the cell types that are first or predominantly affected (e.g., rod-cone degeneration, cone dystrophy, cone-rod degenerations, etc.), the portion of the retina that is primarily involved (center vs. periphery, such as in Stargardt disease and retinitis pigmentosa, respectively), and/or the presence of disease progression (stationary vs. progressive).^8,9^ They can also be further classified as non-syndromic (affecting only the eye) or syndromic (affecting the eye along with other organs, such as the auditory or renal systems).^10^

Retinitis pigmentosa (RP) is the most prevalent form of IRDs, characterized by the degeneration of rods, primarily, and cones, at a later stage,^11,12^ whereas cone and cone-rod dystrophies (CD/CRD) are characterized by the exclusive or primary loss of cones, respectively.^13,14^ The generally stationary forms of cone disorders, grouped as color vision disorders (CVD), include achromatopsia, blue cone monochromacy, as well as common color blindness.^15^ Similarly, congenital stationary night blindness (CSNB) is the non-progressive form of rod dysfunction.^16^ Macular diseases (MD), in which degeneration is largely restricted to the macula, include the second most prevalent form of IRDs, Stargardt disease, as well as Best disease, Sorsby macular dystrophy, etc.^12,17,18^ Lastly, some other non-syndromic IRDs indirectly affect photoreceptors or involve other retinal cell types, such as optic atrophies (OA), exudative vitreoretinopathies (EVR), etc.^19,20^ The most severe form of non-syndromic IRDs is Leber congenital amaurosis (LCA), characterized by retinal blindness in early infancy.^21^ Syndromic IRDs, though less common, constitute a more heterogeneous group comprising more than 80 described clinical entities. The most prevalent among them are ciliopathies, such as Usher syndrome (USH), Joubert syndrome, Bardet–Biedl syndrome (BBS), and Senior–Løken syndrome (SLS).^12,22,23^ Phenotypic variability among patients with the same IRD subtype can also be extensive, and include differences in age of onset, rate of progression, severity, etc. Establishing a clinical diagnosis can therefore be a challenging task, often requiring a multidisciplinary approach that combines patient and family medical history with specialized diagnostic tests such as visual acuity and perimetry assessments, electroretinogram (ERG), fundus autofluorescence (FAF), and optical coherence tomography (OCT).^24^

Moreover, despite being monogenic conditions, IRDs are genetically highly heterogeneous and display multiple inheritance patterns (autosomal dominant – AD; autosomal recessive – AR; X-linked; mitochondrial).^25^ Indeed, over 350 genes have been linked to retinal phenotypes, with syndromic forms accounting for ∼200 of them and RP alone being associated with ∼80 genes.^23,26^ Given this genetic complexity, Next-Generation Sequencing (NGS) has become a cost- and time-effective method for the simultaneous screening of multiple genes, especially in large study cohorts. However, the current diagnostic rate for patients, reported in the scientific literature, varies between 53% and 76%, based on results from multiple NGS techniques, such as panel sequencing, whole-exome sequencing (WES), and whole-genome sequencing (WGS).^27–33^ This diagnostic gap could be attributed to technical limitations, undiscovered disease-causing genes, or variants in regions not typically covered by targeted sequencing procedures, such as intronic or intergenic areas. However, emerging techniques, like in vitro RNA splicing assays^34^ and long-read sequencing,^35^ are being developed to address these challenges.

Another major hurdle in routine molecular diagnosis of IRDs is the use of incomplete or outdated lists of disease-associated genes,^36^ which hinders the proper design of real or virtual gene panels, an accurate interpretation of sequencing data, and the establishment of reliable genotype-phenotype associations. In this study, we aim to address this problem by providing an updated list of IRD-related genes, obtained from the latest scientific research, databases of human DNA variations, as well as from repositories of gene expression data. This resource, curated by experts in the field, will be continually updated and made available on a dedicated website, ultimately to help researchers and clinicians identify disease-causing variants and support future discoveries and molecular diagnoses.

## Results and Discussion

### Data mining and identification of genes associated with IRDs

Genes associated with IRDs were identified through data mining of public databases and published literature, and were individually curated by at least two independent experts, according to the procedures described in the Materials and Methods. At the end of the selection process, 468 genes (including four loci: *RP17*, *MCDR1*, *MCDR3* and Xq27.1) were retained based on strong evidence of disease association (Figure 1). Another 196 genes were classified as “Candidates”, primarily due to evidence from only a single affected family, and 16 genes were excluded due to insufficient evidence, conflicting data, or definitive proof of non-association with IRDs. (Table S1).

**Figure 1:**
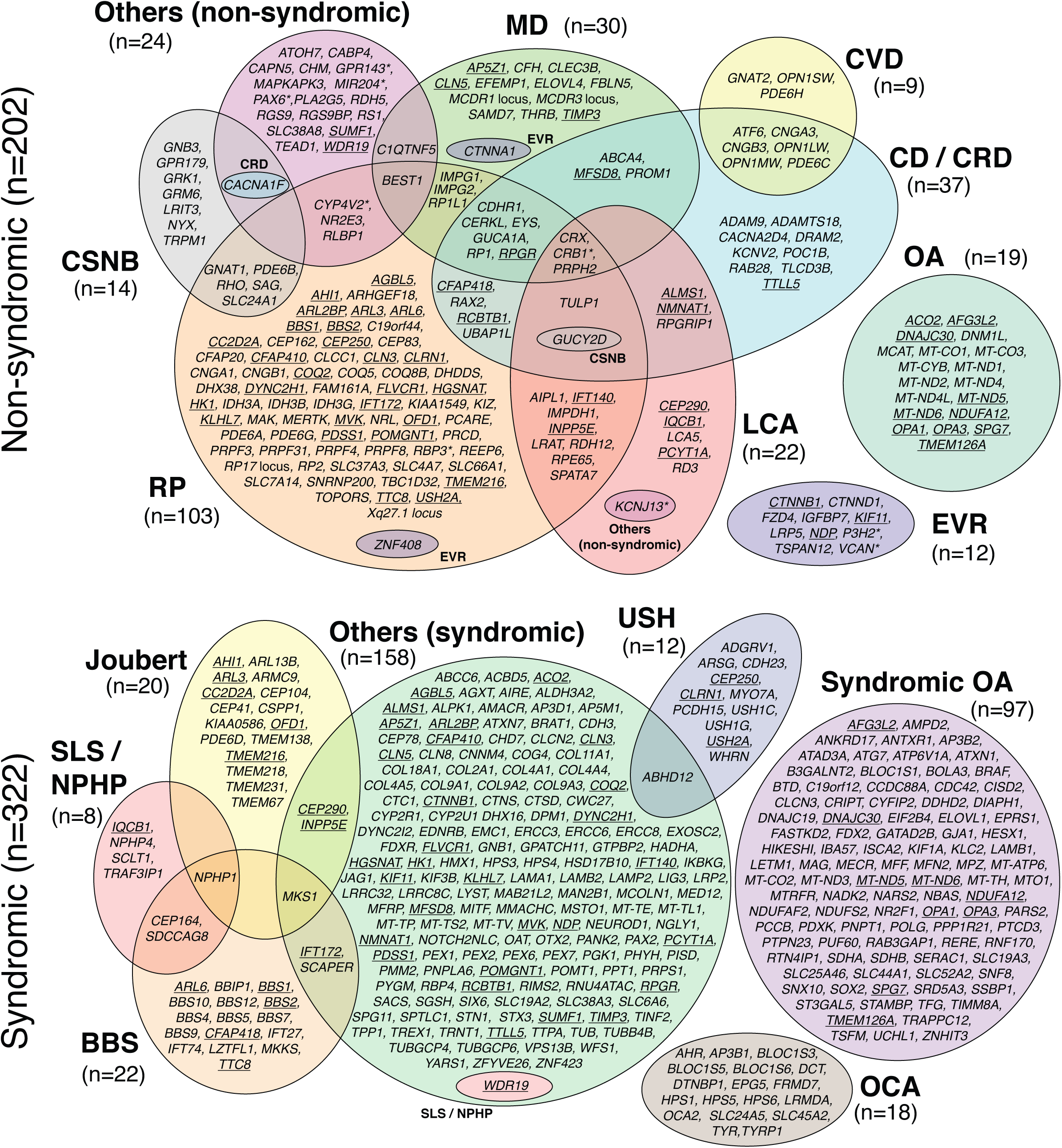
Venn diagram of genes and loci associated with IRDs (total=468). Underlined genes are linked to both non-syndromic and syndromic phenotypes. Asterisks point to genes that can also be involved in non-retinal ocular diseases. n, number of genes.

### Phenotypes and inheritance

We chose a two-level approach to phenotype classification. The first level broadly distinguished between syndromic and non-syndromic phenotypes, based on the presence or absence of multisystemic signs in addition to retinal pathology. The second level defined 16 clinical subsets, including 14 specific groups (e.g. RP, MD, etc.), and two heterogeneous categories that did not fit these groups: “other non-syndromic” and “other syndromic” (Table 1). Notably, we intentionally combined narrowly-defined phenotypes such as Stargardt disease, choroideremia, or Sorsby fundus dystrophy into one of the 14 classes, as these entities are each associated with only one or very few genes (e.g. *ABCA4*, *CHM*, and *TIMP3*, respectively).

**Table 1:** Clinical classification of various inherited retinal diseases (IRDs).

| **Phenotypes and abbreviations** | **Broad category** |
| --- | --- |
| Achromatopsia, color vision abnormalities, color-blindness (CVD) | Non-syndromic |
| Bardet-Biedl syndrome (BBS) | Syndromic |
| Cone dystrophy, cone-rod dystrophy, Stargardt disease (CD / CRD) | Non-syndromic |
| Congenital stationary night blindness (CSNB) | Non-syndromic |
| Exudative vitreoretinopathy, Norrie disease (EVR) | Non-syndromic |
| Joubert syndrome (Joubert) | Syndromic |
| Leber congenital amaurosis (LCA) | Non-syndromic |
| Macular dystrophy (MD) | Non-syndromic |
| Oculocutaneous albinism, foveal hypoplasia (OCA) | Syndromic |
| Optic atrophy, optic nerve hypoplasia (OA) | Non-syndromic |
| Retinitis pigmentosa (RP) | Non-syndromic |
| Senior-Løken syndrome, nephronophthisis (SLS / NPHP) | Syndromic |
| Syndromic optic atrophy (Syndromic OA) | Syndromic |
| Usher syndrome (USH) | Syndromic |
| Others non-syndromic | Non-syndromic |
| Others syndromic | Syndromic |

Out of the 468 curated disease-causing genes and loci, 202 were associated with non-syndromic diseases in 9 phenotypic groups, and 322 were associated with syndromic diseases in 7 phenotypic groups, with 56 genes found to be associated with both non-syndromic and syndromic disease (Figure 1). Overall, most genes had variants responsible for AR inheritance (69%, n=323), followed by AD (15%, n=70), AD-AR (7.7%, n=36), X-linked (4.7%, n=22), and mitochondrial (3.6%, n=17) heredity (Table S1 and Figure 2).

**Figure 2:**
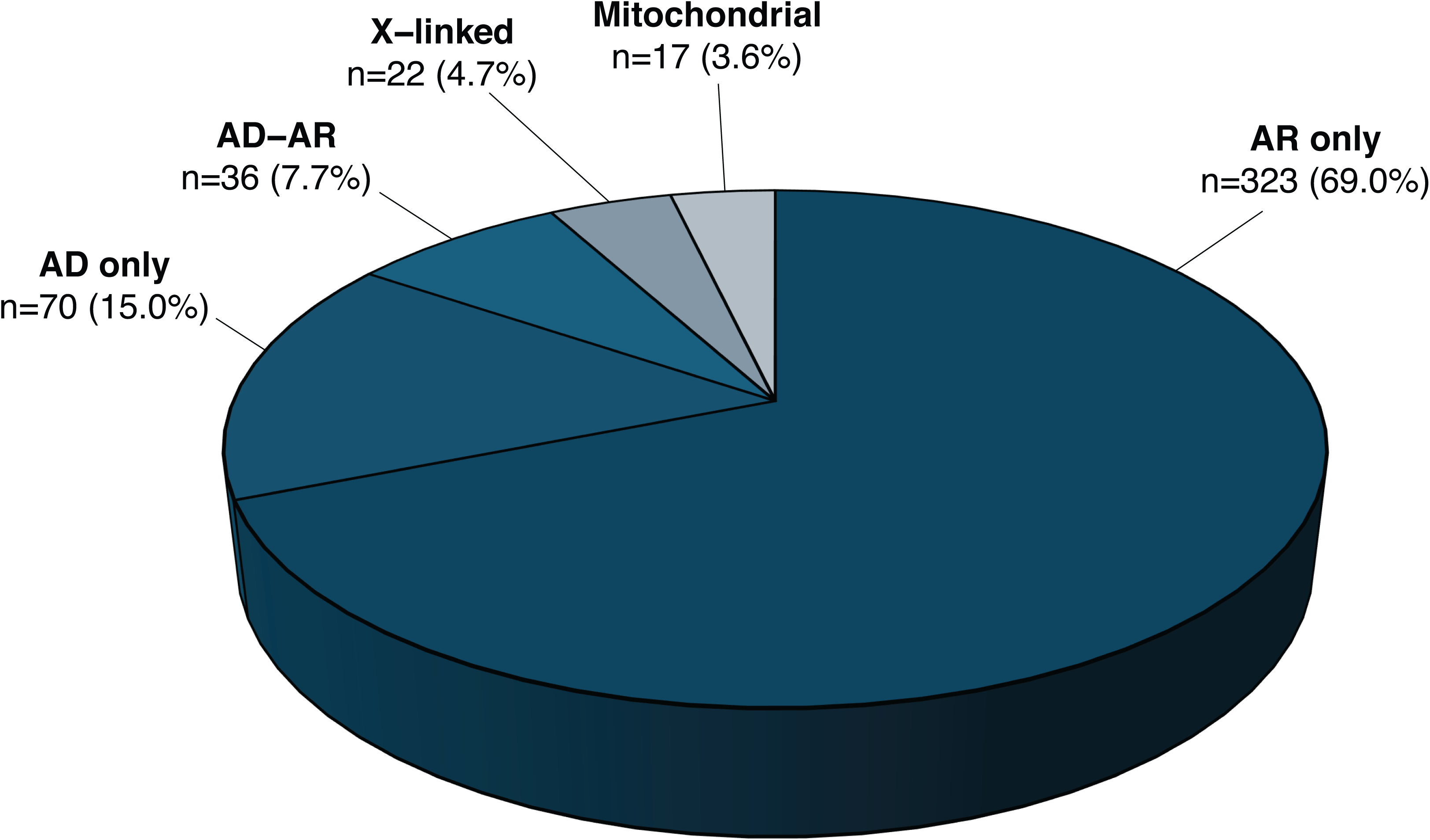
Inheritance mode of diseases associated with all curated genes and loci. n, number of genes.

Among the non-syndromic phenotypes, RP (n=103) involved the highest number of genes, followed by CD/CRD (n=37), MD (n=30), LCA (n=22), OA (n=19), CSNB (n=14), EVR (n=12), and CVD (n=9), while 24 genes were associated with other non-syndromic phenotypes (Figure 1). Notably, within these genes, 52 were associated with more than one phenotype (Table S1, Figure 1). For instance, *CRX*, *CRB1*, and *PRPH2* were each linked to four phenotypes/clinical categories: RP, MD, CD/CRD, and LCA; similarly, *GUCY2D* was associated with RP, CD/CRD, LCA, and CSNB (Figure 1, Figure S1). In several instances, the genotype-phenotype relationship depended on the type of variant [loss-of-function (LoF) vs. missense] or on variant location within specific protein domains.^37–41^ For example, in *CRB1*, loss-of-function variants are typically associated with LCA, while missense variants are more commonly linked to RP or MD.^38,42^ Similarly, disease phenotypes associated with *GUCY2D* vary according to the type and location of the variant. For instance, truncating mutations in the extracellular domain cause LCA, whereas missense changes in the protein kinase domain may result in LCA or CSNB, and variants in other parts of the protein are associated with RP, CSNB, CD/CRD, or LCA.^43^ Likewise, *RPGR* can be linked to RP or CD/CRD depending on the position of the variant along its primary sequence, in a gradient-dependent manner.^44^

For syndromic phenotypes, gene associations included: syndromic OA (n=97), BBS (n=22), Joubert syndrome (n=20), USH (n=12), SLS / NPHP (n=8), oculocutaneous albinism (OCA) / foveal hypoplasia (n=18), and “others (syndromic)” (n=158), a broad and heterogeneous group involving additional systemic involvement beyond the eye (Figure 1B).

Finally, 22 of the 56 genes implicated in both syndromic and non-syndromic conditions encoded ciliary proteins, for which severe variants (typically loss-of-function) tend to cause syndromic forms, while milder changes (typically missense or splicing variants) are more often associated with non-syndromic disease (e.g., in *ARL3*, *CEP290*, or *USH2A*).^45,46^

### Historical perspective

Since the first identification of an IRD-associated gene in 1988 (*OAT*, linked to gyrate atrophy), the number of genes implicated in these conditions has increased steadily, with an average rate of ∼13 novel discoveries per year (Figure 3A). However, this growth has not been uniform. Until 2010, when gene identification primarily relied on linkage analysis, homozygosity mapping or candidate gene approaches,^47^ the trend was essentially linear, with ∼11 new genes identified annually. In 2010, *FAM161A* became the first IRD gene to be identified using next-generation sequencing (NGS), ^48^ marking the beginning of a more rapid phase of gene discovery, which peaked at ∼26 genes per year and lasted until 2018. After 2018, however, the discovery rate declined to ∼8 genes per year, with only 6 genes identified in 2024, despite continued access to high-throughput sequencing. This slowdown may reflect the increasing rarity of newly identified genes in terms of gene-specific genetic prevalence --that is, variants in these genes tend to account for a smaller number of affected individuals in the population (Figure 3B).

**Figure 3:**
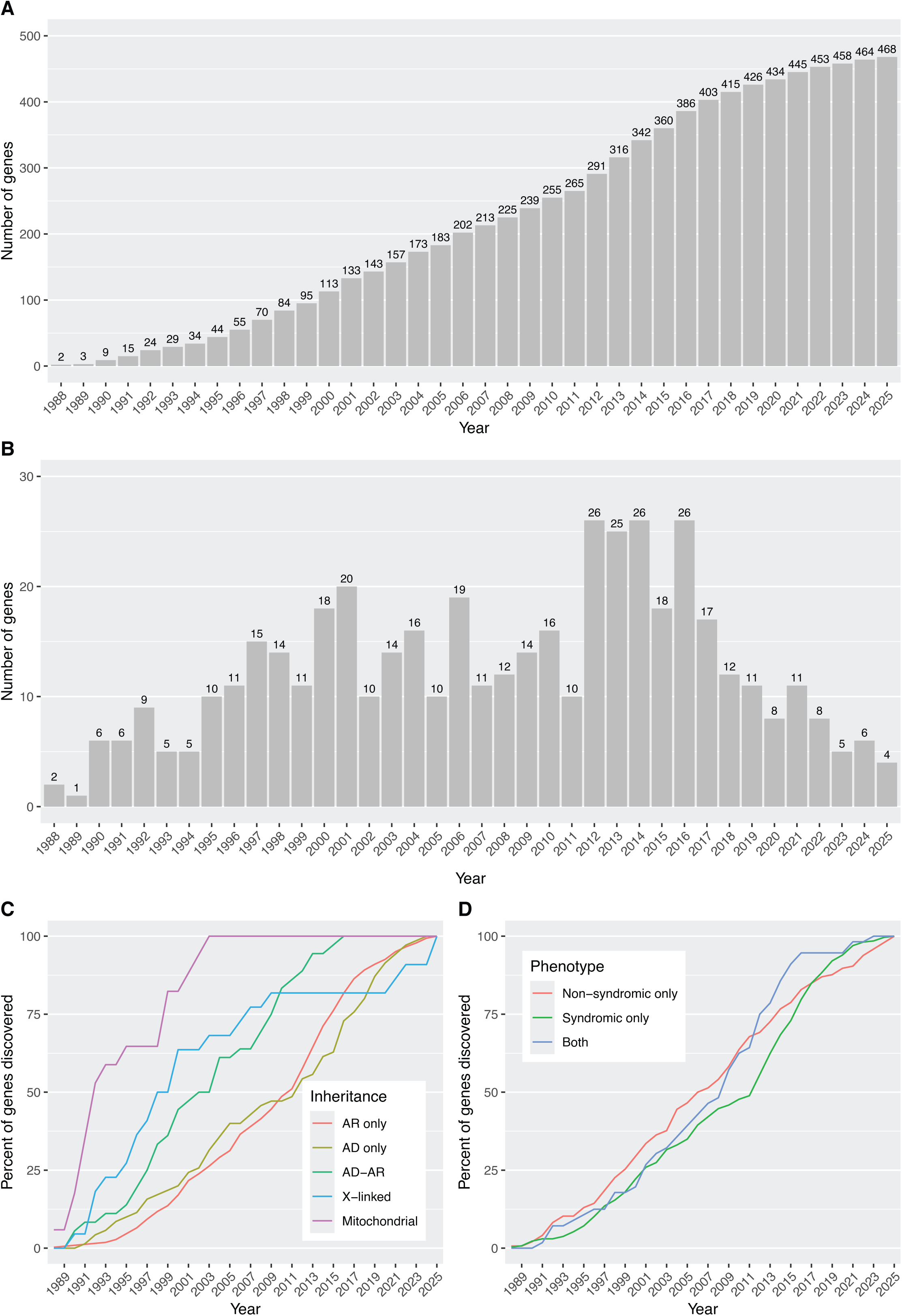
Discovery of IRD genes through time. (A) Cumulative number of genes identified, per year. (B) Annual count of new gene discoveries. (C) Cumulative percentage of genes discovered, stratified by inheritance mode. (D) Cumulative percentage of genes discovered, stratified by broad phenotypic categories.

When stratifying IRD-associated genes by inheritance mode, we observed that those linked to phenotypes due to mitochondrial DNA defects were usually identified in the earliest years, followed by X-linked genes (Figure 3C), despite mitochondrial and X-linked forms representing the least common inheritance patterns in IRDs (Figure 2). This is likely due to the relative ease of detecting mitochondrial and X-linked inheritance, especially in large pedigrees. In addition, balanced X-autosome translocations in affected females and (micro)deletions in affected males facilitated the positional cloning of several X-linked IRD genes (*CHM, NDP, RPGR, RP2*).^49–52^ Similarly, although AD inheritance accounts for only 15.5% of all detected IRD genes (Figure 2), AD phenotypes also tended to be discovered earlier than AR phenotypes. This likely reflects the greater statistical power of linkage analysis in AD families compared to AR families of equivalent size (Figure 3C). In contrast, the rate of gene discovery was relatively similar over time between genes causing non-syndromic and syndromic IRDs (Figure 3D).

### Functional classification

We investigated the biological functions of the 464 curated genes (excluding the 4 loci) by stratifying them into 20 functional categories based on Gene Ontology (GO) terms, as detailed in the Materials and Methods. Genes not assignable to any of these categories were manually reviewed and placed in an “Others” group. In total, 299 genes (64.4%) fell into a single functional category, while 165 genes (35.6%) were classified into multiple categories (Figure 4). These overlapping classifications enabled the identification of functionally related clusters.

**Figure 4.**
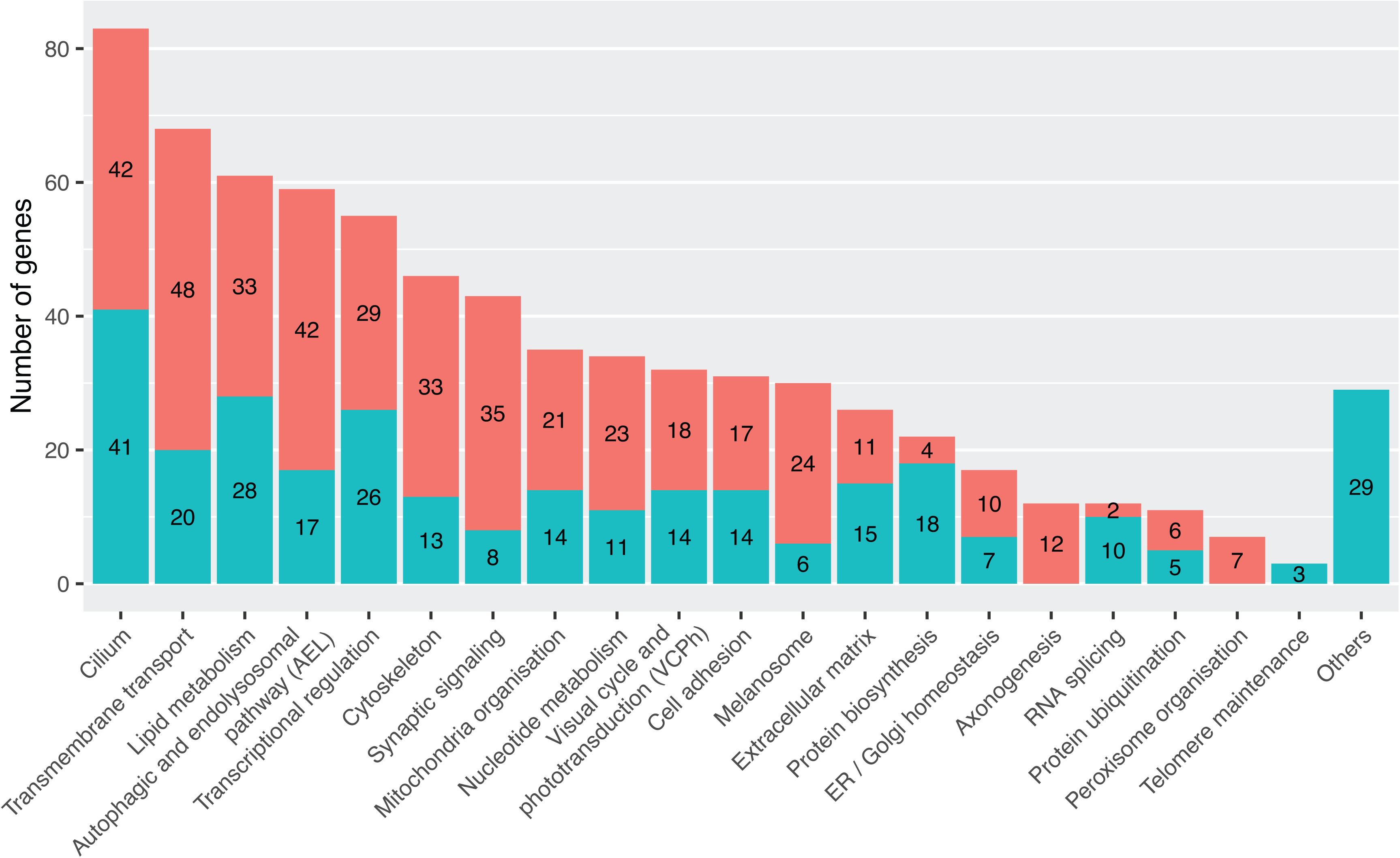
Functional categorization of IRD-associated genes. The bar graph shows the number of genes assigned to each functional category relevant to IRDs. Bar segments indicate whether each gene is annotated in a unique category (blue) or appears in multiple categories (coral). The "Others" group includes genes with roles that could not be confidently assigned to the main categories.

For instance, “Cilium” and “Cytoskeleton” shared 20 genes, reflecting the cytoskeleton’s role as both a structural component of cilia and a regulator of ciliogenesis.^53^ “Cilium” also shared 8 genes with “Melanosome”, and “Cytoskeleton” shared 7 with “Synaptic signaling”, due to the involvement of ciliary structures in melanosome transport and actin filaments in synaptic architecture.^54–58^ Similarly, “Mitochondria” and “Nucleotide metabolism” overlapped by 10 genes, as several rate-limiting steps of nucleotide metabolism occur in mitochondria.^59^ Each of these categories also shared 12 to 16 genes with “Transmembrane transport”, related to mitochondrial electron transport processes.^60^ “Lipid metabolism” and “Visual cycle and phototransduction” (VCPh) shared 9 genes through retinoid metabolism pathways.^61^ Furthermore, 10 to 11 “Lipid metabolism” genes overlapped with “Autophagic and endolysosomal pathway” (AEL) and “Transmembrane transport”. Lastly, AEL, “Synaptic signaling”, “Transmembrane transport”, and “Melanosome” shared 7 to 14 genes, presumably by virtue of their involvement in photoprotection, heterophagy, autophagy in the RPE, and synaptic signal transmission^62,63^ (Figure S2).

“Cilium” was the largest functional category, comprising 83 of the 464 genes (17.9%) (Figure 4). This was expected, given the critical role of cilia in photoreceptor physiology.^64^ The second and third largest categories were “Transmembrane transport” (68 genes, 14.7%) and “Lipid metabolism” (61 genes, 13.2%), encompassing proteins not only involved in VCPh but also in membrane-related metabolism that contributes to IRD pathogenesis. Notably, VCPh ranked tenth, representing only 32 genes (6.9%), reflecting a historical research shift: earlier gene identification efforts specifically targeted VCPh and retina-specific pathways, whereas modern studies are more unbiased.

We next evaluated the distribution of functional categories across three phenotype classes and 16 clinical subsets. In non-syndromic IRDs, the top categories were “Cilium” (37 genes), “Transmembrane transport” (32 genes), and VCPh (30 genes) (Figure S3A). VCPh was particularly retina-specific: 30 of its 32 genes (93.8%) caused non-syndromic IRDs, while other functional classes showed broader phenotypic associations, with less than 54.0% of their genes confined to non-syndromic cases.

“Cilium” and VCPh genes were most commonly associated with RP, while “Transmembrane transport” genes were linked equally to RP and OA (Figure S3B). OA-associated genes also belonged to “Mitochondria” and “Nucleotide metabolism” categories, which primarily contribute to OA. CSNB was mainly associated with “Synaptic signaling” and VCPh genes were consistent with defects in the phototransduction cascade and ribbon synapses. EVR was predominantly linked to “Extracellular matrix”, “Cell adhesion”, and “Transcriptional regulation” genes. Although VCPh genes represented only 30 of the 202 non-syndromic IRD genes, they were associated with the broadest range of clinical subtypes --including LCA, MD, CVD, and CRD (Figure S3B).

In syndromic IRDs, “Cilium” again dominated (68 genes), reflecting the multi-system involvement typical of ciliopathies (Figure S3A). Usher syndrome, Joubert syndrome, SLS, and BBS were mainly linked to “Cilium” and “Cytoskeleton” categories (Figure S3C). OCA was represented majorly by “Melanosome” genes, highlighting the dual role of melanin pathways in ocular and cutaneous pigmentation.^65^ Syndromic OA was associated with nearly all categories, consistent with its complex etiology. In contrast, VCPh was underrepresented in syndromic IRDs, with only two implicated genes—both involved in multiple pathways: *GNB1*, which causes a neurological phenotype,^66^ and *RBP4*, associated with skin involvement^67^ (Figure S3C).

### Inheritance of disease and variant classes

Details about inheritance and phenotypes are shown in Figures 2 and 5A. The majority of curated genes were associated with an AR inheritance pattern, across syndromic and non-syndromic phenotypes. AR genes were more frequently linked to syndromic presentations (n=205) than to non-syndromic forms (n=78). AD inheritance was slightly more common in genes associated with syndromic conditions (n=40) than with non-syndromic ones (n=28), though the difference was modest and AD inheritance was not observed for prevalent conditions such as Usher syndrome, Joubert syndrome, Bardet–Biedl syndrome, etc (Figure S4). In contrast, the discrepancy was more pronounced among genes exhibiting both AD and AR inheritance, with 25 linked to non-syndromic diseases versus only 4 associated with syndromic conditions. X-linked IRDs were evenly represented across syndromic and non-syndromic forms, as well as specific clinical subtypes. Mitochondrial inheritance was observed in both phenotype classes but was almost exclusively associated with OA. This condition results from the degeneration of RGCs, which transmit visual signals to the brain via the optic nerve and rely on high levels of ATP, a process critically dependent on intact mitochondrial function.^19^ Details on inheritance of syndromic vs. non-syndromic phenotypes are shown in Figure S4.

**Figure 5:**
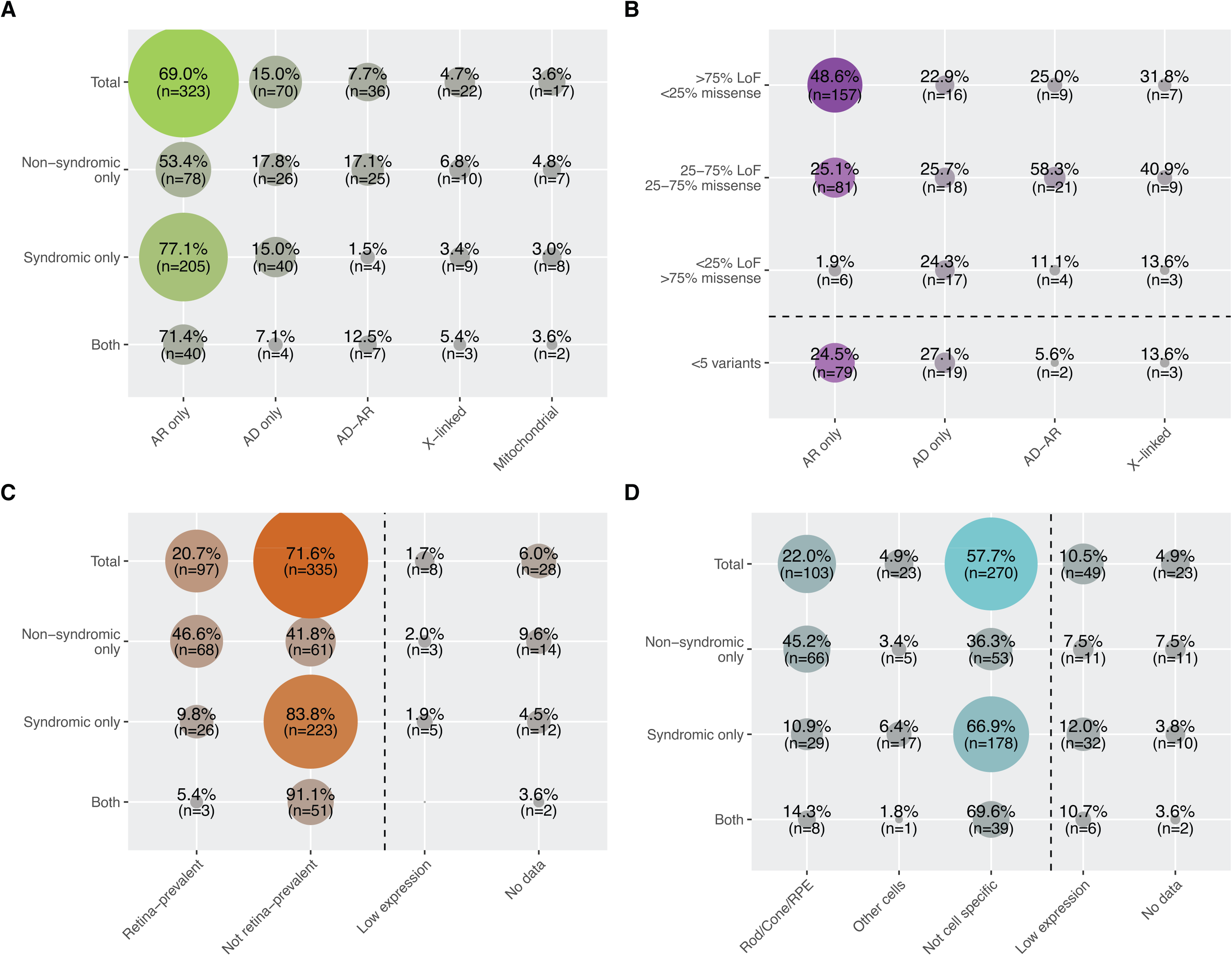
Inheritance and expression features of IRD genes. Co-occurrence matrices between: (A) Inheritance mode and broad phenotypic categories, (B) Inheritance mode and type of pathogenic variants, (C) Expression in retina / other tissues and broad phenotypic categories, (D) Expression in retinal cell types and broad phenotypic categories. n, number of genes.

Next, we investigated whether the spectrum of pathogenic and likely pathogenic (PLP) variants correlates with inheritance mode or phenotypes. Each IRD gene was assigned to one of four categories based on ClinVar^68^ data: (i) >75% LoF variants; (ii) 25–75% LoF and missense variants (mixed); (iii) >75% missense variants; or (iv) fewer than 5 PLP variants (rare). We observed a strong correlation between variant spectrum and inheritance patterns. Genes with AR inheritance were mostly enriched for LoF variants, followed by mixed or rare classes (Figure 5B). This is consistent with the haplosufficiency of most recessive alleles and the fact that AR conditions typically result from complete protein loss. By contrast, AD conditions generally result from heterozygous gain-of-function, dominant-negative, or haploinsufficient mutations. Accordingly, genes in the AD group were nearly evenly distributed across all four variant categories.^69^ Likewise, genes with AD-AR or X-linked inheritance, neither strictly dominant nor recessive, were most commonly found in the mixed LoF/missense group (Figures 5B). LoF variants were also the most prevalent type of DNA changes overall, and mostly associated with syndromic conditions and ubiquitously-expressed genes (Figure S5).

### Gene expression

To investigate a potential correlation between tissue-specific gene expression and the syndromic or non-syndromic nature of retinal disease, we analyzed bulk RNA-seq data from various tissues using the FANTOM5 dataset.^70^ Based on gene expression levels in retinal versus non-retinal tissues, we defined four categories: “Retina-prevalent” and “Not retina- prevalent” for genes with reliable expression patterns across all tissues, and “Low retinal expression” or “No data” for genes with very low expression in the retina or absent from the FANTOM5 dataset (see Materials and Methods).

We found that 46.6% (n=68) of genes causing non-syndromic IRDs were classified as “Retina- prevalent,” which aligns well with classical mechanisms of pathogenesis. Conversely, 41.8% (n=61) of genes associated with diseases restricted to the retina were ubiquitously expressed across tissues (Figure 5C). This is not surprising, as many housekeeping genes, essential for all cells, have been previously implicated in non-syndromic IRDs. These include splicing factor genes and genes involved in core metabolic pathways such as the TCA cycle, Coenzyme Q biosynthesis, and nucleotide metabolism.^71–74^ A widely accepted hypothesis for this paradox is the retina’s intrinsic sensitivity to even minimal metabolic or functional disturbances, making it particularly vulnerable compared to other tissues or organs. Supporting this, the majority of genes causing syndromic IRDs (83.8%, n=223) or involved in both syndromic and non-syndromic forms (91.1%, n=51) were associated with “Not retina prevalent”, many of which are linked to ciliopathies (Figure 5C).

We also examined cell-specific gene expression within the retina in relation to disease phenotype. Using single-cell RNA-seq data, we first classified IRD genes into five categories: “Rod/Cone/RPE,” “Other cells,” “Not cell specific,” “Low expression,” and “No data” (Figure 5D). Notably, 45.2% (n=66) of genes linked to non-syndromic IRDs showed specific expression in photoreceptors or the RPE, compared to 36.3% falling into the “Not cell specific” group. This supports the established concept that non-syndromic IRDs often result from dysfunction or degeneration of cones, rods, RPE cells, or combinations thereof.^75^ This observation is reinforced by the fact that most syndromic IRD genes (66.9%, n=178) showed broad expression across multiple retinal cell types, reflecting their functional relevance in other organs as well. Interestingly, over 15.4% of IRD genes had either minimal or undetectable expression in retinal cells. In addition to developmental stage–specific expression, this is likely due to technical limitations of scRNAseq, particularly transcript dropout events, rather than true biological absence.^76^

We then analyzed scRNAseq expression patterns in the context of specific clinical phenotypes (Figures 6, S6). As expected from clinical and electrophysiological studies, RP was linked to genes expressed in photoreceptors and RPE cells. Rod-specific gene involvement matches the typical clinical course: initial night blindness followed by progressive rod and cone degeneration, culminating in tunnel vision. Macular dystrophies and cone or cone-rod dystrophies could not be reliably distinguished based on scRNAseq alone, likely due to overlapping expression profiles in the same cell types and differences primarily in the affected retinal regions (macula versus the entire retina). Interestingly, CD/CRD showed strong involvement of genes expressed exclusively in cones, consistent with cone-only disease mechanisms. LCA, a condition characterized by early-onset, severe vision loss, involved genes affecting rods, cones, and/or the RPE (but not restricted to a single photoreceptor type).^77^ This suggests that LCA results from disruptions to fundamental cellular processes necessary for the function of all photoreceptors and the RPE. In CVD, gene expression was limited to cones, which is consistent with the fact that color vision depends entirely on this photoreceptor subtype. CSNB is associated with nyctalopia from birth and is typically caused by mutations in genes involved in synaptic junctions between photoreceptors (primarily rods) and bipolar cells, although some CSNB subtypes may also involve milder cone dysfunction, as indicated by our analysis (Figures 6, S6).

**Figure 6:**
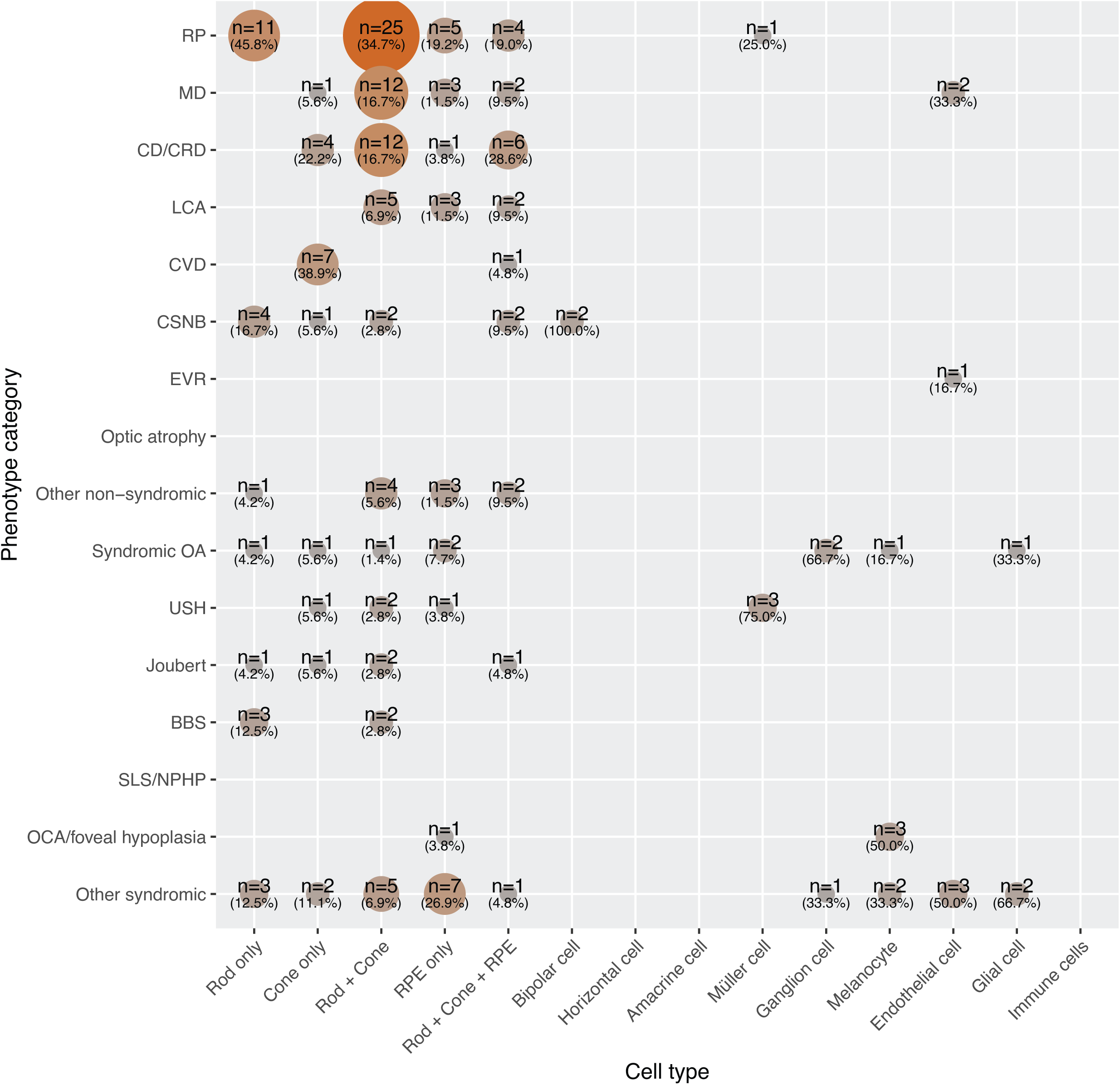
Co-occurrence matrix between phenotypes and retinal single-cell gene expression data. n, number of genes.

The scRNAseq expression patterns for syndromic IRDs were more diverse. For instance, genes implicated in OCA and foveal hypoplasia were expressed exclusively in melanocytes, consistent with their role in melanin biosynthesis. Conversely, some genes associated with syndromic OA were expressed in melanocytes, retinal ganglion, and glial cells, while three Usher syndrome genes showed expression in Müller cells. Notably, *USH1G*, *CLRN1*, and *CDH23*, linked to Usher syndrome, have been shown to be involved in the disease through Müller cell dysfunction as well.^78–80^ Finally, the group of “other syndromic” phenotypes involved genes expressed in a wide range of retinal cell types. This broad expression likely reflects real biological relevance rather than an artifact of phenotype grouping. In contrast, genes associated with non-syndromic conditions (although more numerous overall) tended to show restricted expression limited to photoreceptors and the RPE (Figure 6, top part).

### Implementation of the database

All the data presented here is hosted on a website with an intuitive user interface, ensuring easy navigation and accessibility (https://retigene.erdc.info/). The content will be regularly updated through user requests to reflect the latest information, maintaining both accuracy and relevance for users.

### Conclusions

In summary, we have assembled a freely accessible online database of genes involved in IRDs, which will be continuously updated. This comprehensive catalog is intended to support the identification of disease-causing variants, the development of more accurate genetic testing panels, and a deeper understanding of the molecular mechanisms underlying these conditions. We believe that this resource will contribute to the development of new targeted therapies, improve diagnostic precision, and ultimately enhance patient care for individuals affected by IRDs.

## Materials and Methods

### Operational definition

In this study, we define IRDs as conditions that affect the retina directly, such as diseases caused by mutations in *RHO* leading to rod photoreceptor death,^81^ *CNGA3* leading to non-functional cones,^82^ or *OPA1* leading to ganglion cell death.^83^ We also consider conditions that involve the retina secondarily due to pathology originating elsewhere, such as mutations in *FZD4*, which cause retinal detachment as a consequence of abnormal retinal vascularization,^84^ or in *ABCC6*, which is mainly expressed in the liver and leads to abnormal calcium accumulation in Bruch’s membrane and its subsequent breakage.^85^

### Gene curation

The initial list of genes associated with IRDs was compiled from multiple sources: OMIM^86^ (accessed on 16^th^ January, 2025), ClinGen^87^ (Retina GCEP, accessed on 31^st^ December, 2024), LOVD^88^ (as of 12^th^ May, 2023), Genomics England PanelApp^89^ (Version 7.0), RetNet^90^ (accessed on 16^th^ January, 2025), as well as reviews from the literature.^8,23,91–93^ As of June 1^st^, 2025, data collection was concluded, resulting in a preliminary list of 680 genes and loci.

Each gene was independently evaluated by two experts and included in the downstream analysis if it met either of the following criteria: (i) it harbored distinct pathogenic variants in two or more unrelated individuals or families showing consistent disease phenotype and inheritance pattern; or (ii) it carried the same variant in at least two unrelated individuals or families, supported by strong functional evidence of pathogenicity. Genes that did not fulfill these criteria were classified as “Candidates.” Genes for which conflicting evidence or definitive proof of non-association with IRDs existed were excluded. Loci identified from linkage and association studies without known causative variant(s) were excluded. If they had known causative variants, then they were checked for the criteria of inclusion as stated above. For example, RP17 is a known locus associated with AD-RP due to the presence of complex structural variants which result in ectopic expression of *GDPD1*.^94^ The RP17 locus was therefore retained since these variants segregated in more than 20 families, thus meeting our criteria of inclusion (i).

### Historical perspective selection criteria

Curated genes were annotated with the year of the publication that first linked pathogenic variant(s) in these genes with any form of IRDs.

### Functional classification of genes

The 464 curated genes were annotated for biological process GO terms (GOTERM_BP_FAT) using the Functional Annotation tool from the Database for Annotation, Visualization, and Integrated Discovery (DAVID) knowledgebase (version 2021).^95^ A total of 440 genes were annotated with over 2500 GO terms in total. Based on these annotations, we grouped the genes into 20 functional categories, as listed in Table S2. Lastly, genes that lacked GO term annotations that fitted into these 20 categories were individually assessed by literature review and either manually assigned to one or more of them or grouped into the category “Others”. Corresponding PubMed IDs (PMIDs) for these manual annotations are also provided in Table S2.

Manual annotation was also performed for genes that required refinement beyond the DAVID output due to an unspecific GO term assigned. For example, *PDE6B* and *PDE6C* were directly grouped into “visual cycle and phototransduction” functional category based on the GO terms listed in Table S2, while other related genes—*PDE6A, PDE6G, and PDE6H*—were annotated under the more generic term “visual perception” (GO:0007601), necessitating their reclassification for consistency across functionally similar genes.

### Tissue and retinal cell type expression specificity

To investigate tissue expression of the 468 curated genes and loci in the human transcriptome, the FANTOM5 RNA expression dataset^70^ was downloaded from The Human Protein Atlas.^96^ The dataset contains normalized transcript-per-million (nTPM) values for 18,287 genes in 60 different human tissue samples, including the retina. The average nTPM values of genes were calculated by grouping some of the tissues as follows: “brain_max” = amygdala, caudate, cerebellum, thalamus, hippocampus, nucleus accumbens, temporal cortex, pituitary gland, putamen, postcentral gyrus, spinal cord, substantia nigra, corpus callosum, frontal lobe, insular cortex, olfactory bulb, pons, occipital pole, occipital lobe, occipital cortex, medulla oblongata, medial temporal gyrus, medial frontal gyrus; “glands_max” = salivary gland, thyroid gland, thymus; “digestive_max” = colon, esophagus, small intestine, appendix, gallbladder, smooth muscle; “heart” = heart muscle; “liver”; “lung”; “pancreas”; “muscle” = skeletal muscle; “lymph_node” = lymph node; “diversive_max” = prostate, spleen, tongue, urinary bladder, adipose tissue, breast, cervix, endometrium, ovary, vagina, placenta, seminal vesicle. Values from testis, kidney, and retina were not grouped. At the end of this process, tissue samples were organized into 13 distinct sets for downstream analysis.

Then for each gene, the z-score_retina_ (nTPM_retina_ vs nTPM of other tissues) and the expression ratio (ratio_retina_, nTPM_retina_ / nTPM_max_ in other tissues) were calculated using a custom R script. Based on these values, genes were further categorized as follows: “Retina prevalent” = z- score_retina_ > 3 AND ratio_retina_ > 3 AND nTPM_retina_ > 1; “Not retina prevalent” = z-score ≤ 3 AND ratior_etina_ ≤ 3 AND nTPM_retina_ > 1; “Low expression” = nTPM_retina_ ≤ 1; “No data” = genes that are not included in the FANTOM5 dataset.

Similarly, for the investigation of the single-cell expression of the 468 curated genes within the human retinal tissue, library-normalized transcripts per cell of the adult human peripheral retina were downloaded from a public repository.^97^ This dataset contains expression normalized to 10,000 transcript counts per cell type for 57118 genes and 53 cell/cell subtypes of the retina. The 53 cell and cell subtypes were condensed into 19 major groups (Rods, Cones, RPE, Horizontal cells, Amacrine cells, Bipolar Cells, Ganglion cells, Muller cells, Astrocytes, Glial cells, Choroidal melanocyte, Microglial, Monocytes, NK cells, T cells, Mast cells, Pericytes, Fibroblasts, Vascular endothelial cells) by taking the average expression of the cell subtypes. For example, group ‘Cones’ is the average of the L/M and S cone sub-cell types. Further, 5 broader groups were created from these 19 major groups namely, “Rods+Cones” (average of the major groups Rods and Cones), “Rods+Cones+RPE” (average of the major groups Rods, Cones and RPE), “Endothelial cell” (average of major groups Pericytes, Fibroblasts, and Vascular endothelial cells), “Immune cells” (average of major groups NK cells, T cells, and Mast cells), and “Glial cells” (average of major groups Astrocytes, glial cells and microglial cells).

Then, for each gene, the z-score (normalized expression count of each group vs average of the normalized expression count of remaining groups) and the expression ratio (normalized expression count of each group divided by the average of the normalized expression count of remaining groups) were calculated on a custom R script. Genes were said to be specific to one of the 24 categories (19 major groups and 5 broader groups) based on z-score, ratio, and nTPM as follows: “Group specific” = z-score_group_ > 3 AND ratio_group_ > 3 AND nTPM_group_ ;if they did not meet this criteria for any of the 24 categories, they were marked “Not cell specific”; “Low expression” = nTPM_retina_ ≤ 1; “No data” = genes that are not included in the scRNAseq dataset. For interpretability, we collapsed the results into three overarching specificity categories: “Rods/Cones/RPE” if the gene passed the “Group specific” threshold to be specific to either rods, cones, or RPE, “other cell” if the gene passed the threshold to be specific to either horizontal cells, amacrine cells, bipolar cells, ganglion cell, muller cell, melanocyte, endothelial cell, glial cell or immune cell and “None” if it was found not to be specific to any of the retinal cell groups.

### Variant classification

VCF file containing variant information for all the curated genes was downloaded from the ClinVar database (version of 21.01.2023). PLP variants were selected and annotated using ANNOVAR.^98^ Then, missense and LoF variants were counted per gene. LoF mutations were defined if the variant led to a canonical splicing event, stopgain mutation, or insertion/deletion leading to a frameshift.

## Data and code availability

All curated genes and figures from this study are available on the RetiGene website (https://retigene.erdc.info/).

## Acknowledgments

E.D. was supported by the Ghent University Special Research Fund (BOF22/DOC/229). M.B. and L.V. received funding from the Research Foundation Flanders (1SD8924N to M.B., 11PS324N to L.V.). L.F.C. is supported by the Centro de Investigación Biomédica en Red (CIBER). G.G.G. acknowledges grants from the Instituto de Salud Carlos III (ISCIII) (CP22/00028 and PI22/01371), also co-funded by the European Union, and from the European Union through the HORIZON programme (HORIZON-HLTH-2023-TOOL-05-04, BETTER, 101136262). L.K.H. received support from the Foundation Fighting Blindness Project Program Award (PPA-0622-0841-UCL). C.A. is supported by ISCIII of the Spanish Ministry of Health (PI22/00321), Centro de Investigación Biomédica en Red Enfermedades Raras (CIBERER, 06/07/0036), IIS-FJD BioBank (PT13/0010/0012), the Organización Nacional de Ciegos Españoles (ONCE), the European Regional Development Fund (FEDER), and the University Chair UAM-IIS-FJD of Genomic Medicine. S.B. received funding from Fondazione Telethon (PE00000006, CUP H93C22000660006-MNESYS). F.C. is supported by the Research Foundation Flanders (G0ACQ24N). F.P.M.C. was supported by the Foundation Fighting Blindness USA (BR-GE-0120-0775-LUMC). C.F.I. and C.T. received funding from RP Fighting Blindness and Fight for Sight UK (RP Genome Project GR586). R.K.K. was supported by The Montreal Children’s Hospital Foundation, The Vision Sciences Research Network (VSRN), The National Institutes of Health (NIH)(R01 EY030499-01, Dr. Lentz), The Canadian Institutes for Health Research (CIHR), Fighting Blindness Canada (FBC), and Fonds de Recherche du Québec - Santé (FRQS). R.K.K. also participates in the NAC Attack clinical trial, which is funded by the National Institutes of Health via grants UG1EY033286, UG1EY033293, UG1EY033286, and UG1EY033292. J.M.M. was supported by ISCIII (PI22/00213, AC21_2/00022, and FORT23/00021; the latter co-funded by the European Union), and by the Generalitat Valenciana (CIPROM/2023/26). C.R. received support from the Swiss National Science Foundation (grant No. 204285).

---

Figure S1: Venn diagram of genes and loci associated with the most common non-syndromic IRDs. Underlined genes are linked to both non-syndromic and syndromic phenotypes. Asterisks point to genes that can also be involved in non-retinal ocular diseases.

Figure S2. Overlap among functional categories associated with IRD genes. Functional categories are listed along both axes. n (or plain numbers), number of genes.

Figure S3: Functional classification of genes, stratified by phenotypes. (A) Broad phenotypic categories. (B) Non-syndromic phenotypes. (C) Syndromic phenotypes.

Figure S4: Co-occurrence matrix between phenotypes and their inheritance. (A) Non-syndromic phenotypes. (B) Syndromic phenotypes. n, number of genes.

Figure S5: Co-occurrence matrices between types of pathogenic variants and (A) broad phenotypic categories or (B) specific tissue expression from bulk RNA-Seq. n, number of genes.

Figure S6: Co-occurrence matrices between phenotypes and scRNAseq data. (A) Retinal- prevalent genes (also minimally expressed in other tissues). (B) Not retinal-prevalent genes, n, number of genes.

