## Supplementary figures and images for "RetiGene, a comprehensive gene atlas for inherited retinal diseases (IRDs)"

### Figure S1

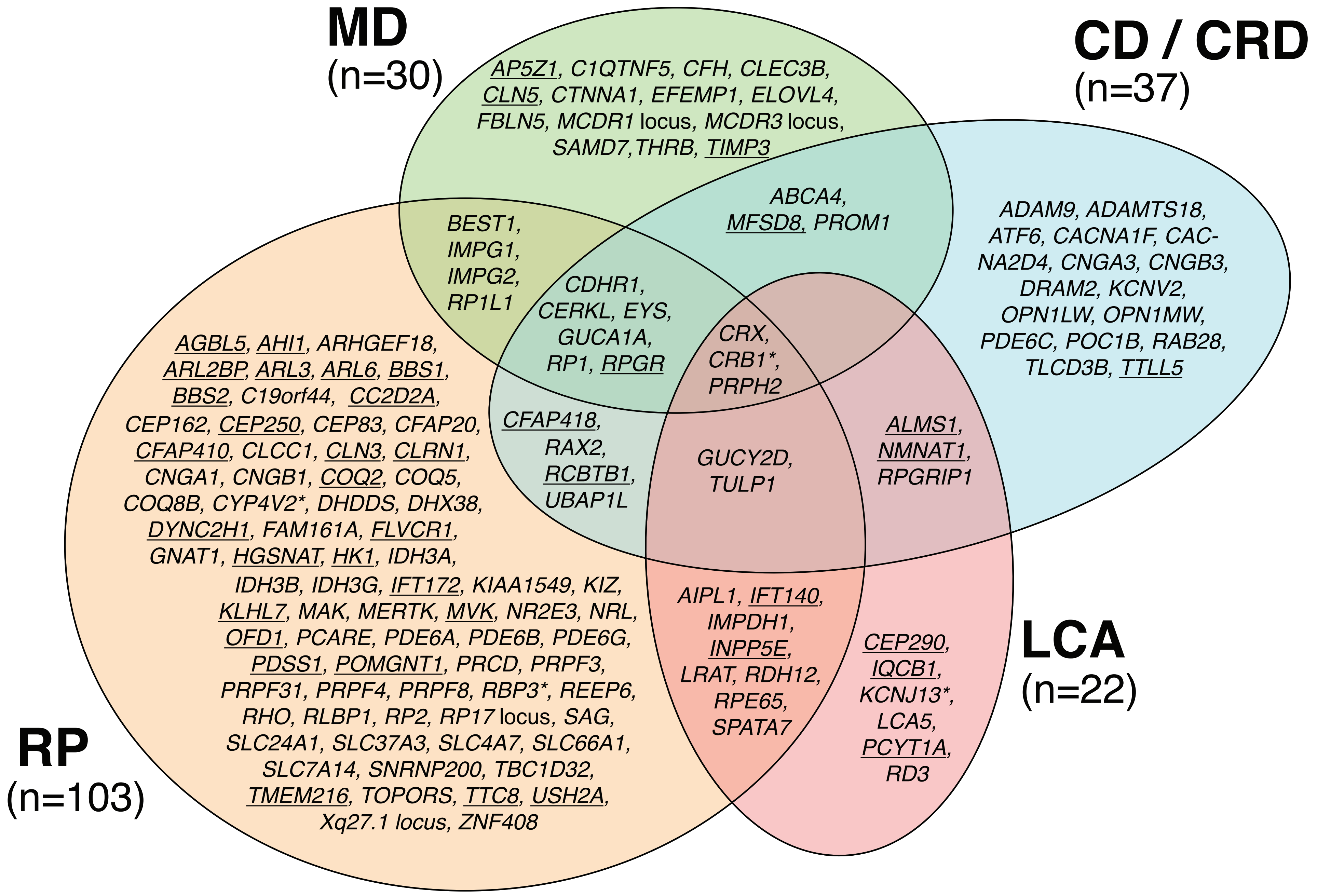


Supplementary Figure 1

### Figure S2

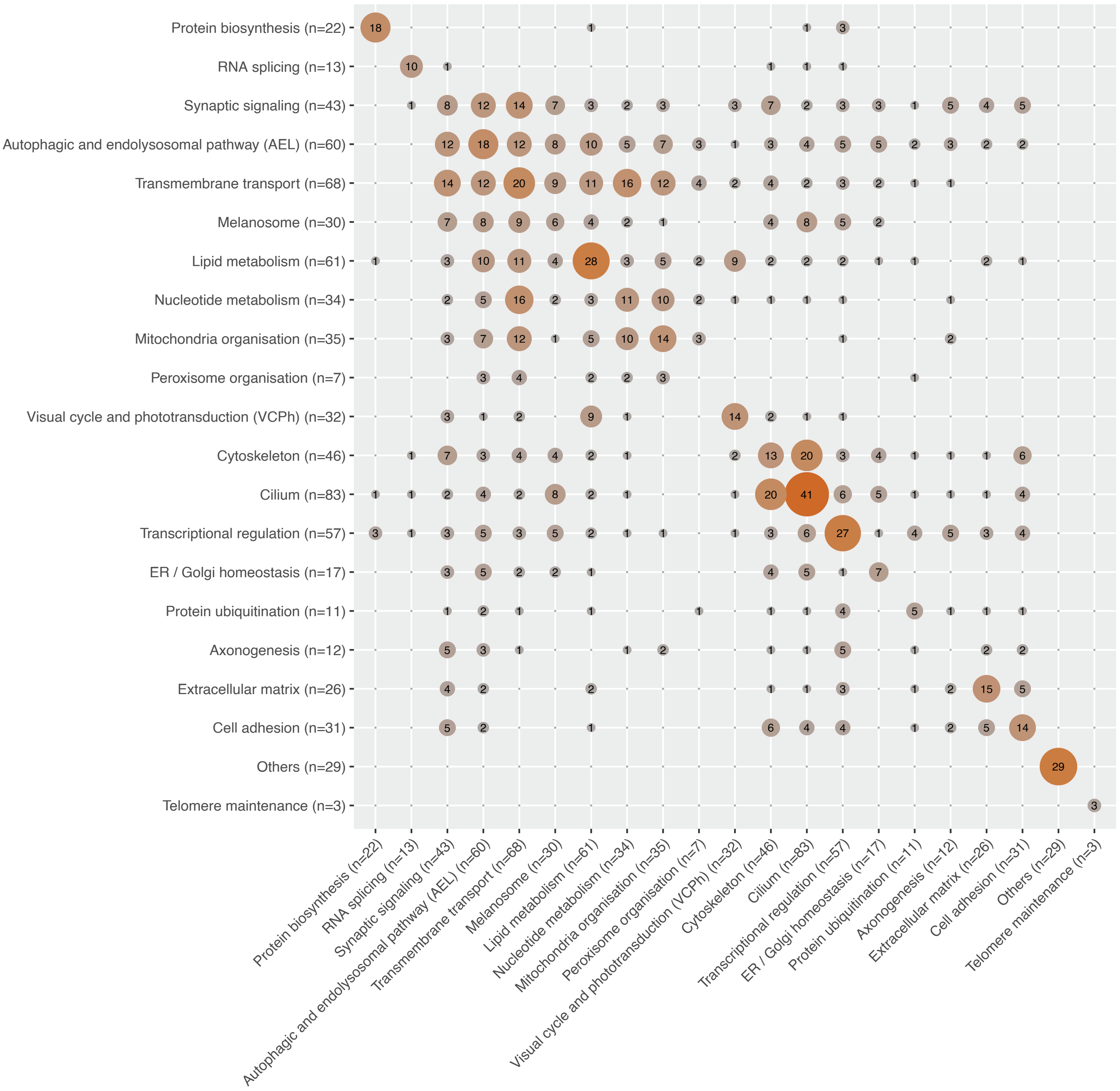


Supplementary Figure 2

### Figure S3

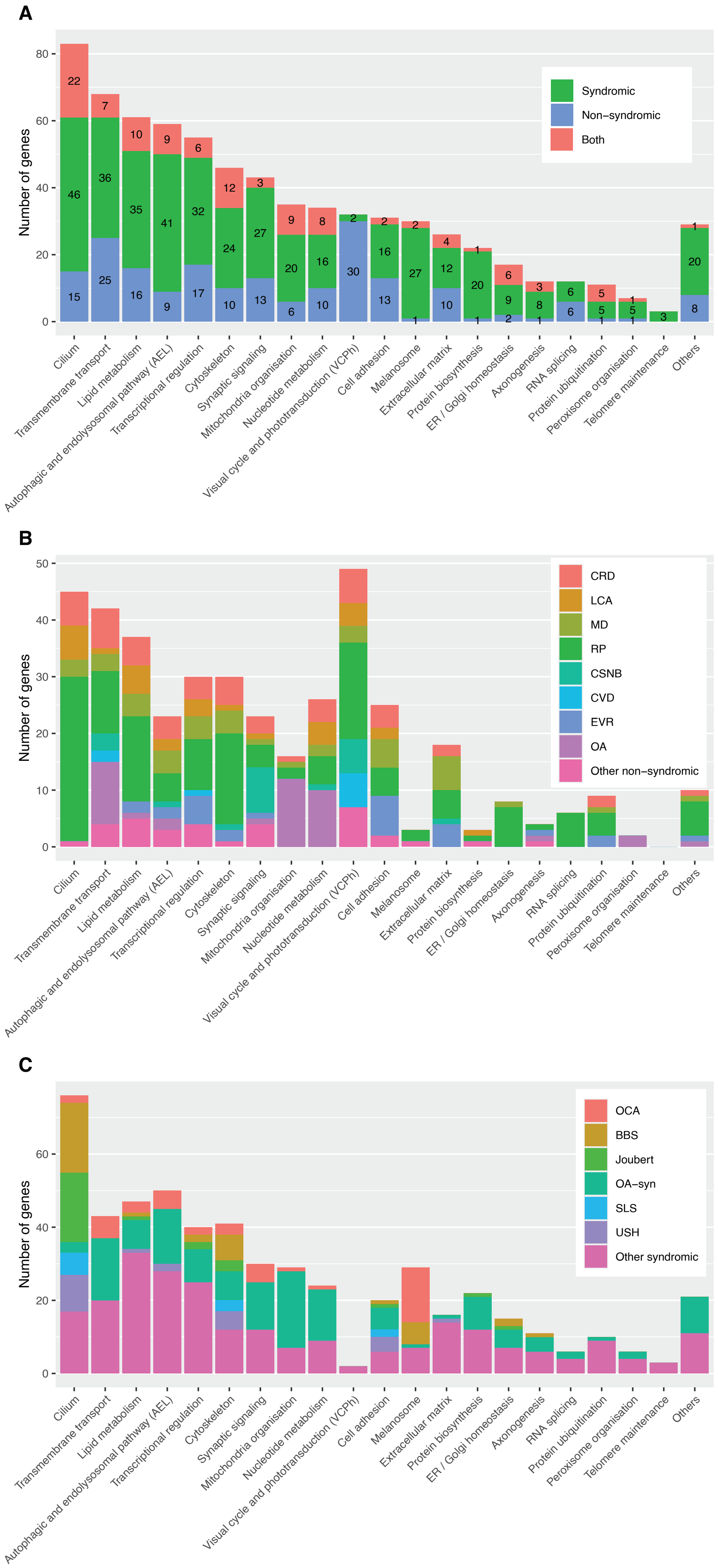


Supplementary Figure 3

### Figure S4

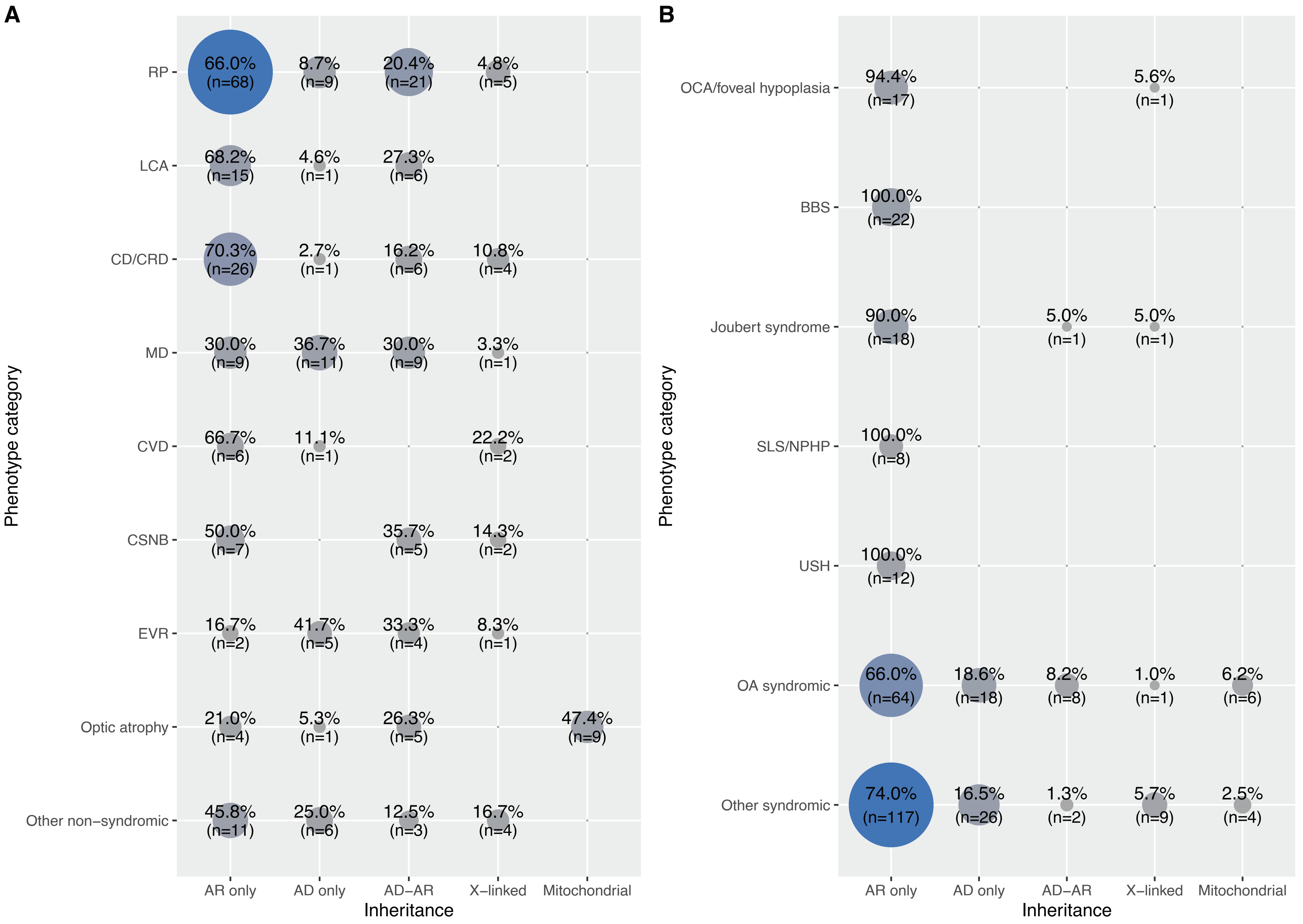


Supplementary Figure 4

### Figure S5

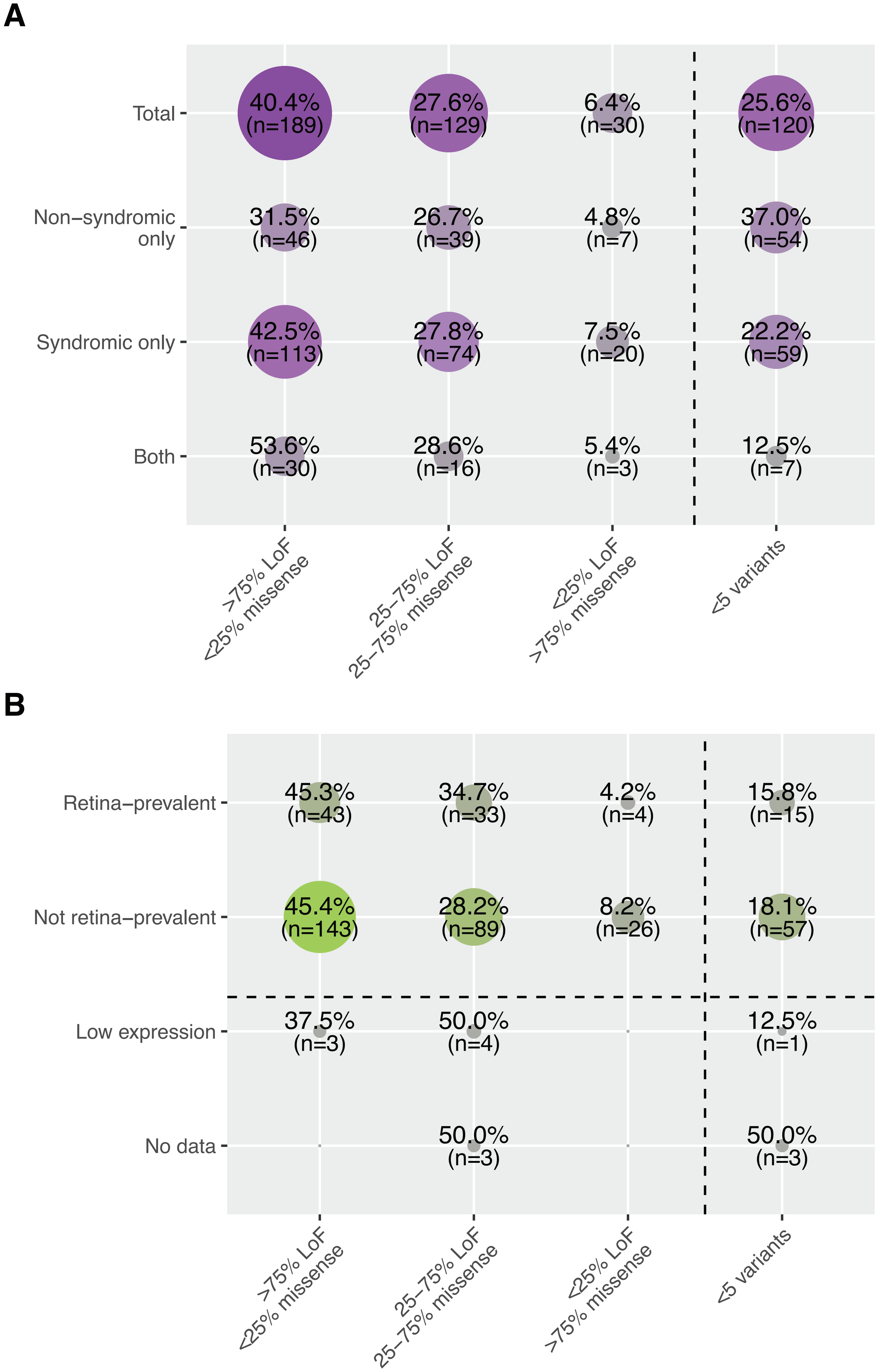


Supplementary Figure 5

### Figure S6

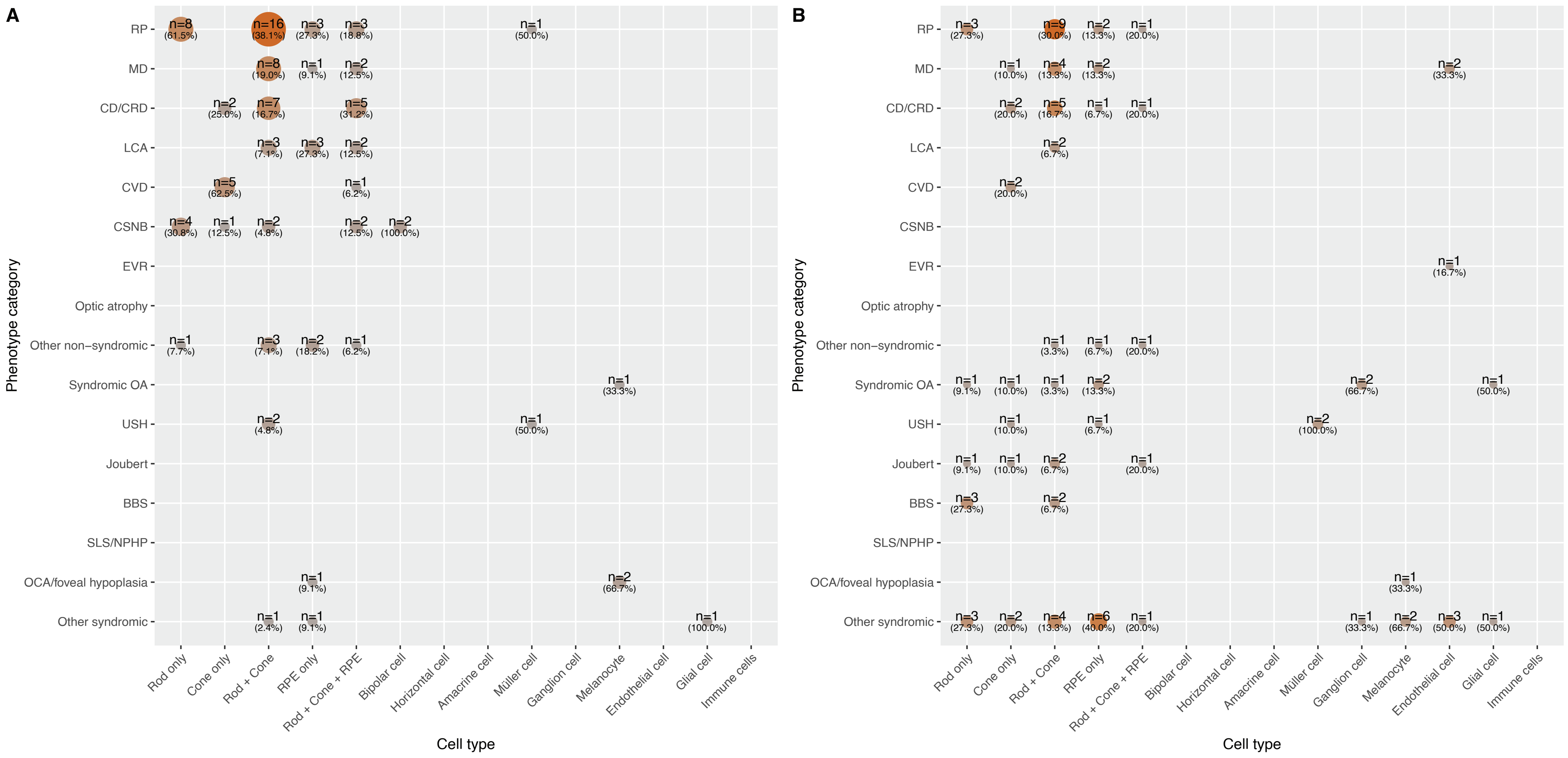


Supplementary Figure 6
